# Abnormal shear stress induces ferroptosis in endothelial cells via KLF6 downregulation

**DOI:** 10.1101/2025.09.10.675441

**Authors:** Jingang Cui, Zhiyu Fan, Suoqi Ding, Jiazhen Zhang, Huihong Shen, Syeda Armana Zaidi, Yongsheng Ding

## Abstract

Stable laminar flow maintains vascular tone regulation, while abnormal blood flow, such as disturbed flow or extreme shear stress, causes endothelial dysfunction, but the underlying mechanism is yet to be explored. We used a microfluidic device to deform flat microchannel into tunnel-like macrochannel. The cross-sectional area of this macrochannel varies with the flow rate and the thickness of the deformable layer, creating three levels of shear stress: low (0.99 dyn/cm^2^, LSS), medium (4.78 dyn/cm^2^, MSS), and high (24 dyn/cm^2^, HSS). Comparing different shear stress exposure to endothelial cells for 24 h, prominent ferroptosis features emerged under either LSS or HSS compared to MSS. These features included increased C11 BODIPY-labeled lipid peroxidation and 4-hydroxynonenal accumulation, CoQ10 depletion, reduced SLC7A11 protein expression, and diminished cell death with Ferrostatin-1 treatment. RNA-seq analysis (LSS/MSS) showed that LSS significantly downregulated transcription of cholesterol homeostasis and unfolded protein response (UPR). Compared to MSS, Western blot results showed that both LSS and HSS reduced the expression of two key enzymes (MVD and IDI1) in the mevalonate pathway, as well as the expression of two main UPR signaling regulators (PERK and BiP). Based on the binding prediction between transcription factors and gene promoters from differentially expressed genes identified through RNA-seq, we found KLF6 to be a key transcription factor. It regulates the PERK-mediated UPR and the mevalonate pathway, which are associated with SLC7A11 expression and CoQ10 synthesis, respectively. The overexpression of KLF6 restored SLC7A11 and CoQ10 levels under both LSS and HSS, significantly reducing foam cell formation, monocyte adhesion, and lipid peroxidation. Our findings reveal KLF6 as a crucial regulator of atherosclerosis induced by abnormal shear stress.

## Introduction

Atherosclerosis is the common pathological basis of a variety of cardiovascular diseases, such as coronary heart disease, myocardial infarction, and cerebral infarction. It is characterized by the accumulation of cholesterol lipid substances in the arterial walls, forming plaques that lead to the narrowing of blood vessels. It is well recognized that the early stage of plaque formation involves endothelial cell (EC) damage, monocyte adhesion, macrophage differentiation, oxidized low-density lipoprotein (oxLDL) phagocytosis, then foam cell formation.

The primary risk factors for atherosclerosis in the circulatory system are well-known: hypertension, hyperlipidemia, and hyperglycemia. A substantial body of evidence from both animal and human specimens suggests that atherosclerosis tends to develop at the bends and branches of arteries [1]. This spatial preference is believed to be linked to changes in blood flow in these regions, marked by low shear stress or oscillatory flow [2-8]. In response to shear stress, several mechanically sensitive transcription factors associated with atherosclerosis have been identified. These include anti-atherosclerotic factors Kruppel-like Factors KLF2 [9-11] and KLF4 [9, 12-14], MEF2[15], and NRF2 [16], as well as pro-atherosclerotic factors AP-1 [17], NF-κB [18], and YAP [19, 20]. These factors play various roles in inflammation, oxidative stress, and cell proliferation. KLF2 and KLF4 have dual functional domains (transcriptional activation and repression), allowing them to regulate genes bidirectionally. They positively regulate endothelial nitric oxide synthase (eNOS) and negatively regulate vascular cell adhesion molecule-1 (VCAM-1). Unlike KLF2 and 4, KLF6 only has a transcriptional activation domain and has also been reported to be involved in the occurrence of atherosclerosis [21, 22]. Recently, azathioprine for psoriasis and apremilast for hypertension have been reported to alleviate endothelial dysfunction and inflammatory responses by restoring KLF6 expression [23, 24].

Endoplasmic reticulum (ER) stress refers to the accumulation of unfolded or misfolded proteins in the ER in response to stimuli, such as oxidative stress, metabolic disorder, and ischemic injury. The unfolded protein response (UPR) is an adaptive cellular mechanism to alleviate ER stress. The accumulation of unfolded or misfolded proteins in the ER causes the 78 kDa glucose-regulated protein (GRP78/BiP) to dissociate from three ER membrane-bound sensors: protein kinase RNA-like ER kinase (PERK), activating transcription factor 6 (ATF6), and inositol-required enzyme 1 (IRE1), thereby initiating three branches of the UPR signaling cascade. To date, various in vitro models have been used to study the impact of shear stress on the ER in endothelial cells. Initially, Fever et al. applied pro-atherosclerotic shear stress using a cone-and-plate flow device, finding that it upregulates BiP through the activation of p38 and integrin α2β1 [26]. A decade later, Bailey et al. used a fan-shaped microfluidic chamber to apply linearly declining shear stress, demonstrating that low shear stress induces VCAM-1 expression and ER stress [27]. More recently, Kim and Woo used a cone system to apply laminar unidirectional flow (up to 12 dyn/cm^2^), revealing that higher shear stress inhibits ER stress-induced apoptosis via the PI3K/Akt-dependent signaling pathway [28]. Based on these in vitro models, low shear stress at atherosclerosis-susceptible sites seems to trigger ER stress and UPR, fostering inflammation and apoptosis. Conversely, high shear stress protects ECs from ER stress. Moreover, some reviews also highlight in vivo studies that show ER stress in endothelial cells at locations susceptible to atherosclerosis [8, 25]. However, there is still limited understanding of how shear stress influences the molecular mechanisms of ER stress in endothelial cells.

From a physiological point of view, the effect of shear stress should be investigated based on fluids with physiological pressure. None of the aforementioned in vitro models of fluid shear stress have been described with a pressure range comparable to physiological blood flow conditions. Our group has recently developed a microfluidic chip composed of three layers of polydimethylsiloxane (PDMS). This chip features deformed tunnel-like channels with varying cross-sectional areas, allowing for multiple combinations of pressure and shear stress [29]. Herein, we established low, medium, and high shear stress levels at 0.99 (LSS), 4.78 (MSS), and 24 (HSS) dyn/cm^2^, respectively, while keeping the on-site pressure within the physiological range of approximately 70 mmHg. After 24 hours under three different shear stress conditions, HUVECs exposed to both LSS and HSS exhibit significant ferroptosis features compared to those under MSS. RNA sequencing analysis comparing LSS and MSS revealed enrichment in the inhibition of steroid biosynthesis and the UPR pathway. Specially, the genes involved include MVD and IDI1, enzymes in the mevalonate pathway, and BiP and PERK, which are crucial for UPR signaling. These genes are related to ferroptosis through their effects on CoQ10 production and SLC7A11 expression in ECs. KLF6, rather than KLF2 and KLF4, is a crucial transcription factor for ferroptosis induced by LSS and HSS. These findings offer new insights into the mechanisms of shear stress-induced atherosclerosis.

## Results

### LSS and HSS induces ferroptosis in endothelial cells

Blood flow shear stress and pressure are two distinct yet interconnected mechanical forces affecting the blood vessel wall. The ideal research approach involves changing one variable while maintaining the others constant, which helps identify each specific effect on the cells. **Figure 1A** shows the microfluidic chip, emphasizing the different shear stress conditions linked to the cross-sectional dimensions of the fluid bodies. LSS, MSS, and HSS can be attained by modifying the thickness of the middle layer, using the given formula and a fixed number of parameters. With a fixed flow rate of 1.6 ml/min under varying shear stress conditions, the fluid pressure in the tunnel-like channels fluctuates around 70 mmHg. This is based on the calculation formula we previously reported, which primarily depends on the flow rate and the size of the downstream guide channel [29]. **Figure 1B** shows a significantly higher number of propidium iodide (PI)-stained dead cells under both LSS and HSS compared to MSS. This result aligns with the change of extracellular LDH level shown in **Figure 1C**, indicating that both LSS and HSS can lead to significant endothelial cell damage. **Figure 1D-H** highlights key characteristics of ferroptosis, including lipid peroxidation (marked by 4-hydroxynonenal and C11 BODIPY), decreased levels of SLC7A11 and CoQ10, and the prevention of cell death with Fer-1 treatment.

**Figure 1.**
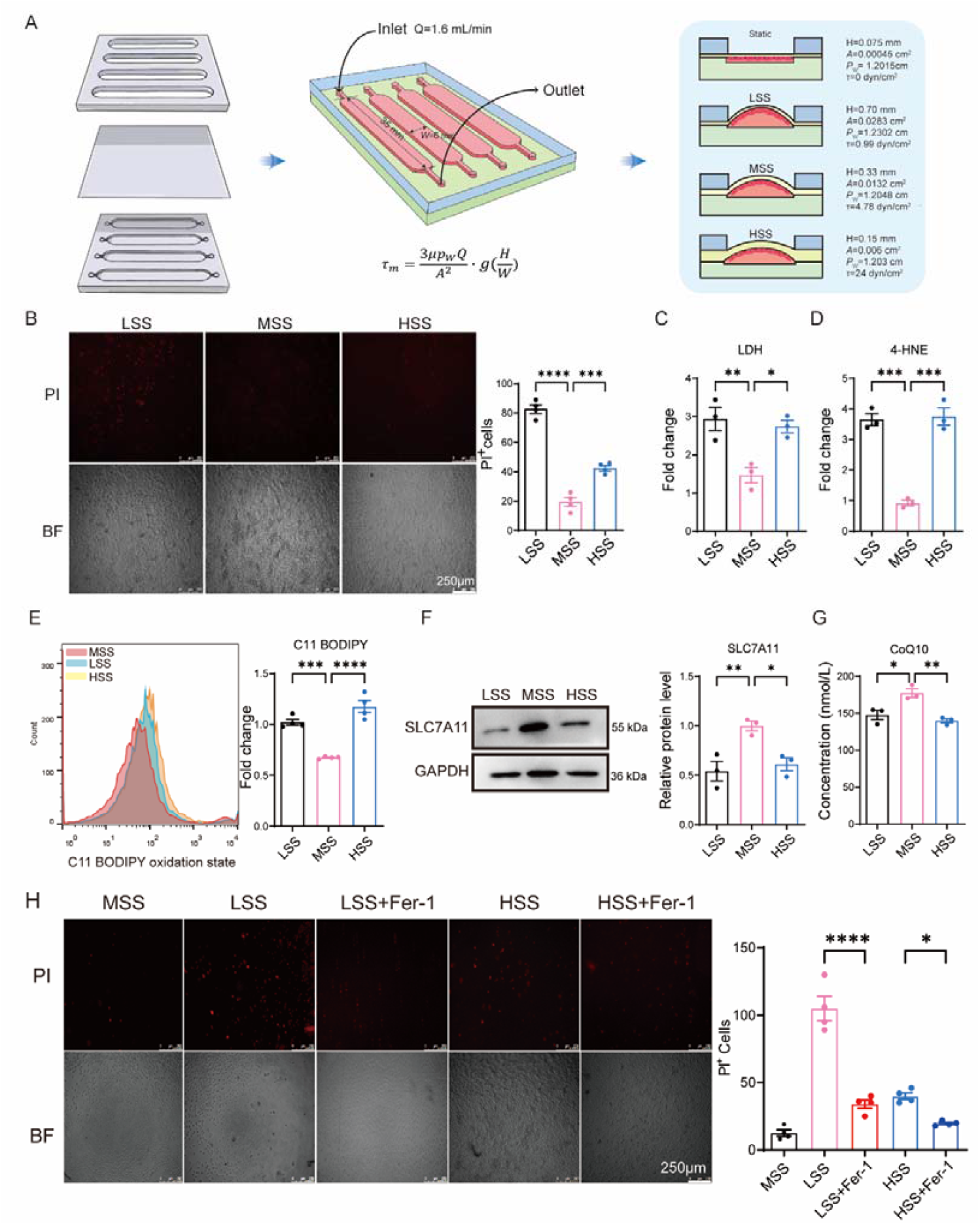
Abnormal shear stress induces endothelial ferroptosis. (A) Diagram of the microfluidic device and calculation of the shear stress. Three levels of shear stress (τ) are formed in the tunnel-like channel, depending on the varying cross-sectional area of flow: LSS = 0.99 dyn/cm^2^, MSS = 4.78 dyn/cm^2^, and HSS = 24 dyn/cm^2^. These values were calculated using a formula that includes the dynamic viscosity of the fluid (*μ*), flow rate (*Q*), channel width (*W*), cross-sectional area (*A*), wetted perimeter (*p*_*w*_), membrane bulge height (*H*), and a function of *H/W (g(H/W)*). (B) Fluorescent microscopic results of PI staining in ECs (The scale bar = 250 μm); (C) ELISA results of extracellular LDH in ECs; (D, E) Flow cytometry results of lipid peroxidation in ECs indicated by 4-HNE and C11 BODIPY. (F, G) Protein expression of SLC7A11 and intracellular CoQ10 level. (H) The results of Fer-1 (10 μM) treatment. Data are expressed as the mean ± standard deviation from three independent experiments (n=3). Statistical significance was determined using one-way analysis of variance (ANOVA) followed by Dunnett’s post hoc test for comparisons among three or more groups, with \**p* < 0.05, \*\**p* < 0.01, \*\*\**p* < 0.001, and \*\*\*\**p* < 0.0001.

### RNA-seq analysis shows that LSS inhibits cholesterol biosynthesis and the UPR

RNA-seq analysis was conducted to systematically explore the mechanisms of endothelial cell damage caused by the abnormal shear stress (LSS/MSS). As shown in **Figure 2A**, LSS significantly downregulated 298 genes and upregulated 100 genes compared to MSS (│log2│> 0.6, Qvalue < 0.05). KEGG, GO, and GSEA shown in **Figure 2B-D** shows a significant enrichment in steroid biosynthesis and ER stress response pathways. Specially, LSS significantly inhibited terpenoid backbone synthesis and UPR. **Figure 2E** indicates that 18 DEGs related to lipid synthesis were significantly downregulated by LSS. Among them, HMGCS1, MVD, and IDI1 are three genes that encode key enzymes in the mevalonate pathway. This pathway is essential for the biosynthesis of sterol isoprenes, such as cholesterol, and non-sterol isoprenes, like ubiquinone. These enzymes catalyze reactions that convert acetyl-CoA to (S)-3-hydroxy-3-methylglutaryl-CoA (HMG-CoA), decarboxylate to isopentenyl diphosphate (IPP), and isomerize between IPP and dimethylallyl diphosphate (DMAPP). These processes are crucial for CoQ10 synthesis, an antioxidant that helps prevent ferroptosis. **Figure 2F** indicates that 11 DEGs related to the UPR were significantly downregulated by LSS. Genes like SDF2L1, HSPA5, HSP90B1, HYOU1, and FICD are involved in protein folding, while HERPUD1 and DERL3 encode proteins for endoplasmic reticulum-associated protein degradation (ERAD). Additionally, PERK and CHOP play roles in UPR signaling. As LSS inhibits these UPR-related genes that manage ER stress, the ability to resolve stress from unfolded or misfolded proteins is likely to be disrupted. To confirm the RNA-seq analysis predictions, we evaluated the protein expression of four genes—MVD, IDI1, PERK, and HSPA5 (BiP)—under various shear stress conditions. As shown in **Figure 2G**, Western blot results indicate that the expression of BiP, PERK, MVD, and IDI1 was significantly reduced by both LSS and HSS compared to MSS. Notably, HSS has a similar effect on the expression of these proteins as LSS, despite the lack of RNA sequencing analysis between HSS and MSS. This suggests an adaptive range of organisms to environmental mechanical stimuli, akin to a ‘Window Effect’.

**Figure 2.**
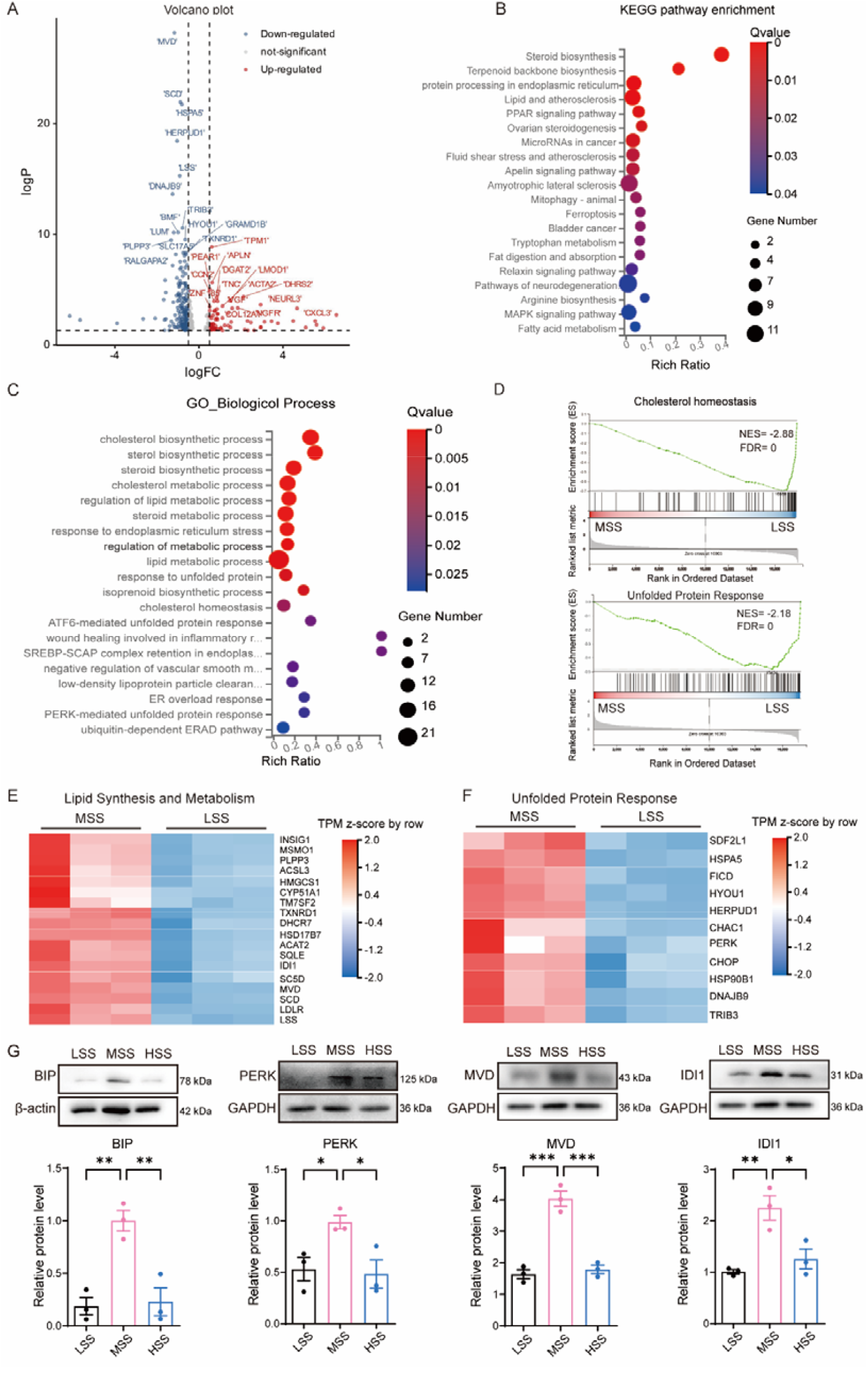
RNA-seq reveals LSS suppresses cholesterol biosynthesis and UPR. (A) Volcano plot of DEGs (LSS vs. MSS;│log2FC│>0.6, Q<0.05); (B) KEGG pathways; (C) GO biological processes; (D) GSEA of steroid biosynthesis and UPR. NES: normalized enrichment score. FDR: false discovery rate. (E) Heatmap of DEGs in lipid synthesis and metabolism. (F) Heatmap of DEGs in UPR. (G) Protein expression of BiP, PERK, MVD, and IDI1 under different shear stress conditions. Data are expressed as the mean ± standard deviation from three independent experiments (n=3). Statistical significance was determined using one-way analysis of variance (ANOVA) followed by Dunnett’s post hoc test for comparisons among three or more groups, with \**p* < 0.05, \*\**p* < 0.01, \*\*\**p* < 0.001, and \*\*\*\**p* < 0.0001. KEGG: Kyoto Encyclopedia of Genes and Genomes, GO: gene ontology, GSEA: gene set enrichment analysis.

### KLF6 is found to regulate the PERK-mediated UPR and the mevalonate pathway

Inhibiting PERK-mediated UPR has been shown to decrease SLC7A11 protein expression in colorectal cancer cells [30]. In our study, both LSS and HSS inhibit this pathway and the mevalonate pathway, impacting SLC7A11 expression and CoQ10 synthesis, which contributes to endothelial ferroptosis. To further understand the impact of shear stress on the UPR and mevalonate pathways, we identified 11 transcription factors with decreased expression under LSS from the downregulated DEGs, including KLF6, MEF2C, ZNF91, and ZNF185, most of which have zinc finger motifs (**Figure 3A**). Using the silicon method, the promoter regions of PERK, HSP5A, MVD, and IDI1 are predicted to contain KLF6-specific binding elements (**Figure 3B**). This is consistent with the Western blot results in **Figure 3C**, showing that both LSS and HSS significantly reduce KLF6 protein expression compared to MSS. To confirm KLF6’s crucial role in regulating these target genes, we created an endothelial cell line overexpressing KLF6 for a gain-of-function experiment (**Figure 3D**). The results in **Figure 3E-F** show that the expression levels of PERK, BiP, MVD, and IDI1 were significantly higher in KLF6-overexpressing endothelial cells compared to the negative control, regardless of whether the cells were treated with LSS or HSS. Additionally, the overexpression of KLF6 largely restored the expression of these four target proteins to levels similar to those observed under MSS.

**Figure 3.**
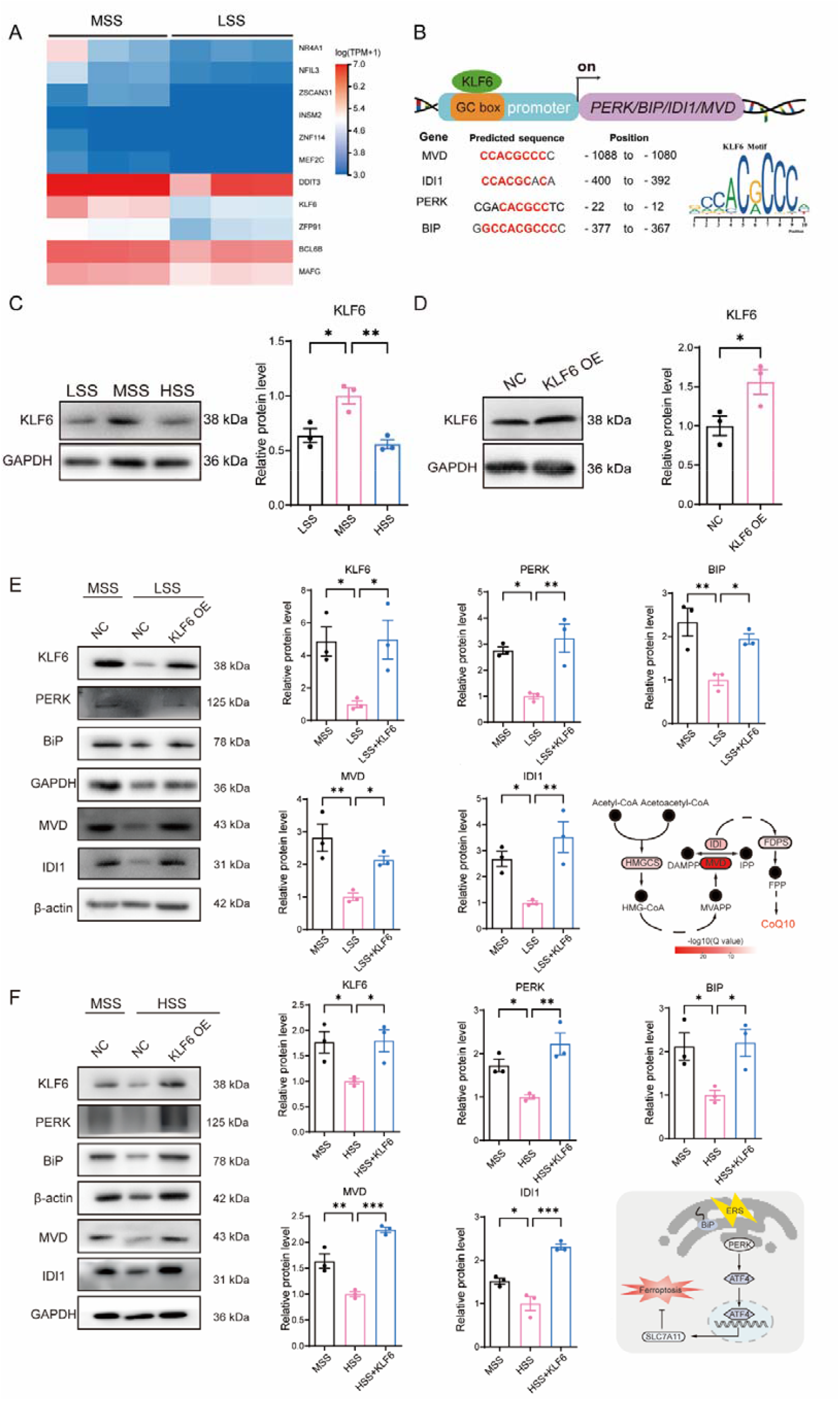
KLF6 regulates the UPR and mevalonate pathway. (A) Heat map of the identified TF genes downregulated by LSS; (B) Predicted KLF6-binding motifs in promoters of PERK, HSPA5, MVD, IDI1 (JASPAR database; matrix ID: MA0472.1); (C) Protein expression of KLF6 under different shear stress conditions; (D) Validation of KLF6 overexpression by Western blot (OE: overexpression, NC: negative control); (E, F) KLF6-OE largely restores the expression of PERK, BiP, MVD, and IDI1 under LSS and HSS to levels seen under MSS. Data are expressed as the mean ± standard deviation from three independent experiments (n=3). For comparisons involving three or more groups, statistical significance was determined using one-way analysis of variance (ANOVA); whereas for comparisons between two groups, an unpaired two-tailed Student’s t-test was employed, with \**p* < 0.05, \*\**p* < 0.01, \*\*\**p* < 0.001, and \*\*\*\**p* < 0.0001.

### Verification of KLF6’s role in endothelial ferroptosis and early atherosclerotic events caused by abnormal shear stress

As illustrated in **Figure 4A-H**, KLF6 overexpression under both LSS and HSS led to an increase in SLC7A11 protein expression and CoQ10 levels. Conversely, it reduced the extracellular LDH level and lipid peroxidation. These results clearly demonstrate that KLF6 overexpression can significantly inhibit ferroptosis induced by LSS and HSS. Based on our previously developed multi-type cell (VSMC/EC/THP-1)in vitro atherosclerosis model [29], the anti-atherosclerotic effect of KLF6 overexpression on the early events of atherosclerosis was examined. As illustrated in Figure 4I-L, both monocyte adhesion and intracellular lipid accumulation (Oil Red O staining) were significantly reduced in KLF6-overexpressed endothelial cells compared to the negative control. These preliminary findings indicate that KLF6 might play a significant role in atherosclerosis, though further validation through animal studies is necessary.

**Figure 4.**
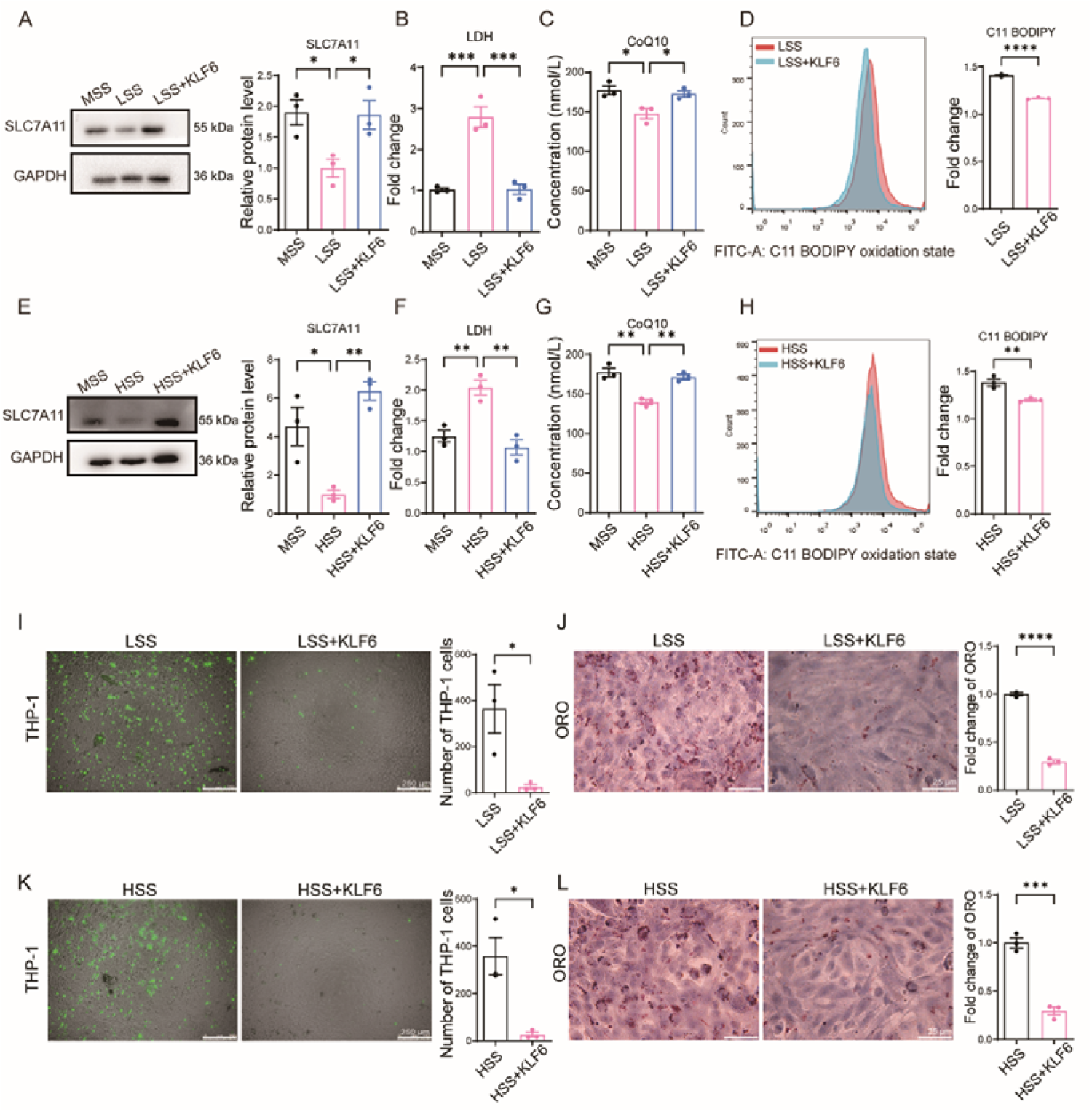
KLF6 overexpression alleviates endothelial ferroptosis and early atherosclerotic events induced by LSS and HSS. (A, E) Protein expression of SLC7A11; (B, F) ELISA results of extracellular LDH; (C, G) ELISA results of intracellular CoQ10; (D, H) Flow cytometry results of lipid peroxidation indicated by C11 BODIPY; (I, K) Fluorescent microscopic results of THP-1 cells (green label) adhering to the VSMC-EC bilayer cell architecture (The scale bar = 250 μm); (J, L) Lipid accumulation visualized by Oil Red O staining (The scale bar = 25 μm). Data are expressed as the mean ± standard deviation from three independent experiments (n=3). For comparisons involving three or more groups, statistical significance was determined using one-way analysis of variance (ANOVA); whereas for comparisons between two groups, an unpaired two-tailed Student’s t-test was employed, with \**p* < 0.05, \*\**p* < 0.01, \*\*\**p* < 0.001, and \*\*\*\**p* < 0.0001.

## Discussion

The endothelium responds to changes in blood flow shear stress caused by the heartbeat, regulating vasodilation and maintaining vascular homeostasis. It is generally believed that the cause of atherosclerosis at the bend or bifurcation of the arteries is closely related to low shear stress or oscillatory shear stress. In vivo studies on the impact of blood flow shear stress on endothelial cells are difficult to perform, so researchers mainly use various in vitro fluid models. However, fluid pressure is frequently overlooked in these studies. Our model demonstrates variations in shear stress at 0.99, 4.78, and 24 dyn/cm^2^, while keeping fluid pressure near the physiological range of about 70 mmHg. To the best of our knowledge, this is the first study to show that LSS and HSS induce ferroptosis in ECs. This result contrasts with the apoptosis observed in other fluid models, possibly due to differences in fluid pressure. Another notable finding is that the EC damage exhibits a V-shaped trend under LSS, MSS, and HSS. This suggests that endothelial cells can adapt to a limited range of shear stress, although the underlying mechanisms are still not understood.

RNA-seq analysis revealed that LSS primarily suppresses the transcriptome of steroid synthesis and UPR-related pathways compared to MSS. Specifically, the transcription of genes such as PERK, BiP, MVD, and IDI1 was reduced, leading to decreased SLC7A11 protein expression and COQ10 synthesis, thereby impairing resistance to lipid peroxidation. Mechanical signal transduction typically involves changes in the conformation of membrane proteins, leading to the depolymerization and remodeling of actin within the cell, activation of transcription factors in the cytoplasm, or epigenetic regulation through histone modifications. Given the complexity of the transduction mechanism for shear mechanical signals, we focus on identifying clues from transcription factors that respond to shear stress. Notably, KLF2 and KLF4, which are shear stress-sensitive transcription factors, are absent from our DEGs dataset, whereas KLF6 is included. Most studies report that KLF6 primarily regulates cell differentiation and tissue development. Additionally, recent research has shown that KLF6 downregulation is also linked to oscillatory flow-induced endothelial cell inflammation and oxLDL-induced endothelial dysfunction [23, 24]. As anticipated, KLF6-overexpressing ECs can restore the BiP-PERK-SLC7A11 and MVD-IDI1-CoQ10 axials under both low and high shear stress conditions. This indicates that KLF6 is crucial for cell survival, consistent with recent findings that KLF6 deficiency significantly exacerbates liver damage, cell apoptosis, and hepatic inflammation [31].

As shear stress rises from low to high, ferroptosis follows a V-shaped pattern, while KLF6 expression displays an inverted V-shape. This suggests that cellular ferroptosis, triggered by both excessive and insufficient shear stress, inversely correlates with KLF6 expression. The results of KLF6 overexpression showed that KLF6 played a significant role in reducing endothelial cell damage and lipid peroxidation under LSS and HSS. In other words, physiologically friendly shear stress promotes KLF6 expression and enhances the antioxidant capacity of endothelial cells. However, the distribution of shear stress in blood vessels is uneven due to variations in vessel diameter and shape, complicating its numerical characterization in vivo. This hinders the assessment of how changes in shear stress impact endothelial cell function in vivo. Consequently, most in vitro models fail to simultaneously replicate the shear stress and pressure present in actual arterial blood flow. A key benefit of our device is its ability to maintain a stable physiological pressure range while delivering a variety of shear stresses. This could explain the ferroptosis we observe, which differs from that in other in vitro models due to abnormal shear stress at a physiological pressure of 70 mmHg. The anti-atherosclerotic effect of KLF6 was also assessed using our previously established atherosclerosis model, which involves a three-cell culture system (SMC, EC, THP-1). This evaluation demonstrated that KLF6 overexpression reduces the adhesion of THP-1 cells and lipid accumulation induced by LSS and HSS.

In summary, we examined the impact of different shear stress on endothelial cells using a microfluidic chip to simulate low, medium, and high shear stress under a physiological pressure of 70 mmHg. Abnormal shear stress levels (0.99 and 24 dyn/cm^2^) significantly induce endothelial ferroptosis by inhibiting the KLF6-UPR/Mevalonate-SLC7A11/CoQ10 dual pathway (**Fig. 5**). Our findings reveal a novel mechanism linking shear stress to the early stages of atherosclerosis, explaining its preference for occurring at arterial bends and branch sites.

**Figure 5.**
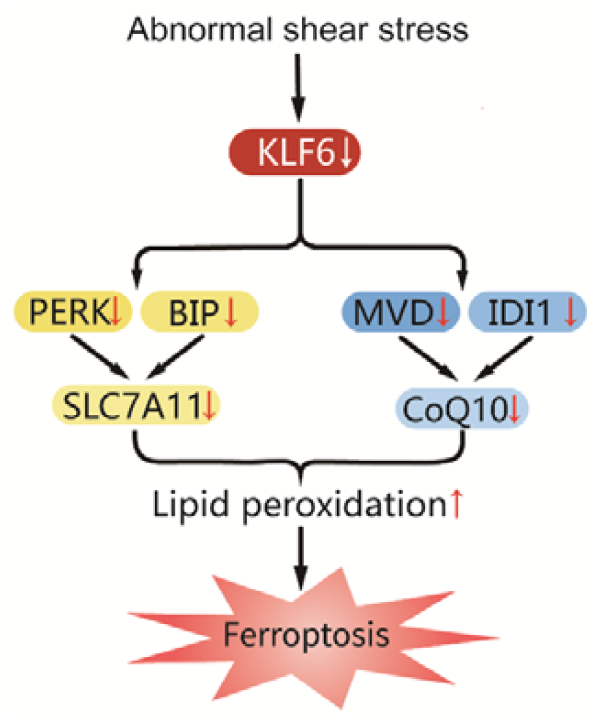
Overview of KLF6 mediated the ferroptosis induced by abnormal shear stress

## Methods and materials

### Antibodies and reagents

Antibodies: SLC7A11 (1:1000, ab175186, Abcam), GRP78/BIP (1:1000, 66574-1-Ig, proteintech), PERK/EIF2AK3 (1:1000, 24390-1-AP, proteintech), KLF6 (1:1000, 14716-1-AP, proteintech), MVD (1:1000, 15331-1-AP proteintech) IDI1 (1:1000, 11166-2-AP, proteintech), Goat Anti-Rabbit IgG-HRP (1:10,000, ASS1006-1, Biorigin), Goat Anti-Mouse IgG-HRP (1:10,000, ASS1007-1, Biorigin). Reagents: Ferrostatin-1 (17729, Cayman), C11-BODIPY581/591 (27086, Cayman), Propidium iodide (P4170, Sigma-Aldrich), Fibronectin (354008, BioCoat), Opti-MEM (31985062, Gibco), M5 HiPer Lipo2000 Transfection Reagent (MF135-01, Mei5bio), TRIzol reagent (15596026, Invitrogen). Dulbecco’s Modified Eagle Medium (DMEM) (12800017, Gibco), Fetal bovine serum (FBS) (10099141, Gibco), LDH detection kit (C0017, Beyotime), Protease inhibitor and phosphatase inhibitor (5872S, Cell Signaling Technology), BCA protein detection kit (P0012S, Beyotime), 4-HNE ELISA kit (RE12203-96T, Bioroyee, China), Oil Red O (ORO) (Biolabs, China), CoQ10 ELISA kit (HBDY-50954H1, Huabodeyi, China) RNAsimple Total RNA Kit (DP419, Tiangen Biotech), CFDA-SE (APExBIO, USA).

### Fabrication of microfluidic chip

The microfluidic chip was made of three PDMS layers. The bottom layer contains multiple parallel shallow grooves (height 0.075 mm, width 6 mm, and length 35 mm). The middle layer is an elastic thin membrane to support cell culture. The top layer is cut with multiple parallel hollow strips for alignment with the shallow grooves of the bottom layer. The hollow strips of the top layer constrain the bulging of the middle layer to form a tunnel-like structure (Figure 1A). The molds used for the top and bottom layers are made of SU-8 negative photoresist embossed on a silicon wafer. The middle layer was obtained by a spin-casted PDMS on a blank silicon wafer. After the middle layer is cured at 65 °C for 15 min, it is contacted by the top layer with the hollowed strips and continued to cure for 1 h to form irreversible binding. Using the plasma oxidation, the embedded groove in the bottom layer is aligned with the hollowed strips in the middle-top layer assembly, and then the two parts are tightly compressed to form a sealed microchannel between the middle and the bottom layers. The assembled microfluidic chip is further cured overnight at 65 °C for use. A series of connecting holes is pre-punched at both ends of the guide microchannel, facilitating its connection to a circulating flow path driven by a peristaltic pump.

### Cell lines and culture

A human aortic vascular smooth muscle cell line (T/G HAVSMC) was obtained from American Type Culture Collection (ATCC, Manassas, VA, USA). A human umbilical vein endothelial cell line (HUVEC-T1) and a human monocytic cell line (THP-1) were purchased from the National Infrastructure of Cell Line Resource of China (Beijing, China). T/G HA-VSMC and HUVEC-T1 were cultured in DMEM (Gibco, USA) containing 10% FBS (Gibco, USA), penicillin (100 U/ml, Gibco, USA), and streptomycin (100 μg/ml, Gibco, USA). Monocytic THP-1 cells were cultured in RPMI 1640 medium (Gibco, USA) supplemented with 10% FBS (Gibco, USA), penicillin (100 U/ml, Gibco, USA), and streptomycin (100 μg/ml, Gibco, USA). The cell culture was maintained at 37 °C in 5% CO2 in a humidified incubator.

### Preparation of cell culture in the microfluidic chip

Before seeding the cells, the channels of the microfluidic chip used for cell culture were coated with a matrix (EHS Matrix, Zhejiang Meisen Zhiyuan Biotechnology Co., Ltd.), which was diluted 100-fold in PBS when used. Inject 300 μL of diluted matrix solution into each channel of the chip. Then incubate the chip at room temperature in the dark for 1 hour to allow coating. After incubation, the coating solution was removed. For experiments applying different shear stresses to HUVECs alone, 300 µL of a HUVEC-T1 suspension with a density of 1.0 × 10^6 cells/mL was injected into the channel. The chip was then inverted and incubated for 24 hours to allow the formation of a stable endothelial cell layer on the middle membrane. On the other hand, for the foam cell formation experiments, 300 µL suspension of HAVSMC with a density of 1.0 × 10^6 cells/mL was first injected into the channels, and 50 µg/mL vitamin C was added to stimulate extracellular matrix secretion. Invert the chip so that the cells settle on the middle membrane and let them continue to culture undisturbed for 24 hours. After that, 300 µL suspension of HUVEC-T1 with a density of 1.0 × 10^6 cells/mL was injected into each channel following the same procedure. After 24 hours, a stable smooth muscle-endothelial cell bilayer structure was formed. Finally, the chip with the firmly established cell layer was connected to a peristaltic pump circuit to apply shear stress. When conducting foam cell formation experiments, 1 mL of THP-1 cell suspension with a density of 2.0 × 10^5 cells/mL needs to be added to the circulating culture medium (15 mL).

### Cell death assay

Cell death was evaluated by propidium iodide (PI) staining and extracellular lactate dehydrogenase (LDH) assay. After 24 h of shear stress treatment, cells were washed three times with PBS and incubated with 5 µg/mL PI in the dark at 37 °C for 30 min. Images were acquired using a fluorescence microscope (DMI6000, Leica). The extracellular LDH release was detected using a commercial LDH detection kit (C0017, Beyotime) and a multi-function microplate reader (Synergy H1 Hybrid Reader, BioTeK) according to the manufacturer’s protocol. To verify the ferroptosis in endothelial cells, the circulating culture medium used during the application of shear stress contained 10 µM Ferrostatin-1.

### Lipid peroxidation measurement

The cell lipid peroxidation was measured by C11-BODIPY 581/591 probe. After 24 h shear stress treated, the cells were washed with PBS three times and incubated with 10 μM C11-BODIPY 581/591 in the dark at 37 °C for 30 min, then the cells were analyzed by flow cytometry.

### ELISA for 4-HNE and CoQ10

The content of 4-HNE in the cells was detected by 4-HNE ELISA kit (RE12203-96T, Bioroyee) according to the manufacturer’s protocol. The content of CoQ10 in the cells was detected by CoQ10 ELISA kit (F13041-A, Beijing Huabodeyi biology science and technology co., ltd) according to the manufacturer’s protocol.

### Western blotting

Cells were treated with shear stress for 24 h, total proteins in the cells were isolated by RIPA lysis buffer (P0013B, Beyotime) with the complete protease inhibitor and phosphatase inhibitor (5872S, Cell Signaling Technology) and protein concentration was determined using a BCA protein detection kit (P0012S, Beyotime). After separation by SDS-PAGE, the proteins were transferred to a PVDF membrane (ISEQ00010, Merck Millipore). The membrane was blocked with 5% non-fat milk in TBST (20 mM Tris, 137 mM NaCl, 0.1% Tween-20, pH7.4) at room temperature for 1 h, incubated with the primary antibodies at 4 °C for overnight, and then the membrane was incubated with anti-mouse or anti-rabbit IgG with HRP linked secondary antibody, followed by chemiluminescence detection (WBKLS0100, Merck Millipore) with a gel imaging analysis system (ChampChemi 910 Plus, Sagecreation).

### RNA-seq data analysis

Total RNA was extracted from the shear stress treated endothelial cells using RNAsimple Total RNA Kit (DP419, TIANGEN BIOTECH) according to the manufacturer’s protocol. RNA quantification and purity were determined using Nanodrop One (Thermo Fisher Scientific). Qualified RNA samples were sent to BGI-Shenzhen (Shenzhen, China) for strand-specific library construction using the NEBNext Ultra II RNA Library Prep Kit (NEB #E7770), followed by sequencing on an Illumina NovaSeq 6000 platform with 150-bp paired-end reads. Raw data processing and downstream analyses were conducted on BGI’s Dr. TOM platform (https://biosys.bgi.com).

### Lentiviral transduction of HUVEC-T1

For the KLF6 overexpression experiment, two lentiviral particles were obtained from GENEWIZ: one containing the human KLF6 gene (plvx-puro-KLF6) and the other an empty plvx-puro vector. These were used for KLF6 overexpression and as a negative control, respectively. In brief, HEK293T cells were transfected with either plvx-puro vector or plvx-puro-KLF6 constructs with psPAX.2 and pMD2.G second-generation lentiviral packaging system. After 48 h incubation, the lentivirus particles in the medium were collected and filtered to infect the HUVEC-T1 cells. After 48 h infection, 4 μg/mL puromycin was added into the cell culture medium to obtain the stable cell lines with successful transduction.

### THP-1 cell adhesion assay

THP-1 cells were incubated with 10 μM CFDA-SE (APExBIO, USA) for 15 minutes at 37 °C. The stained THP-1 cell solution, at a density of 2.0×10^5 cells/ml, was added into the circulating medium. After the cells were treated with shear stress for 24 h, the number of attached THP-1 cells was counted using a fluorescent microscope (Leica, DMI 6000B).

### Oil Red O staining

After the cells were treated with shear stress for 24 h, they were rinsed with PBS three times, and then fixed in 4% paraformaldehyde for 30 min, and then washed again with PBS three times. After the cells were stained with Oil Red O solution (ORO) (Biolabs, China) for 1 h at room temperature, the stained cells were incubated with 60% isopropyl alcohol for 2 min and a hematoxylin solution (Beijing Biodragon Immunotechnologies, China) for 3 min, followed by rinsing the cells with the tap water. The sample was covered with glycerin and then mounted with a slide. ORO-stained cells were photographed under the bright-field condition by a microscope (Leica Microsystems). ImageJ software was used for quantification after setting the appropriate threshold.

### Transcription factor binding site prediction

Putative KLF6-binding sites in promoter regions of PERK (EIF2AK3), HSPA5 (BiP), MVD, and IDI1 were predicted using the JASPAR database (https://jaspar.elixir.no/; version 2022). Genomic sequences (2000 bp upstream of transcription start sites) were analyzed with the KLF6 position weight matrix (MA0472.1). Binding motifs containing GC-box elements (consensus: 5′-CACCC-3′) with a relative profile score threshold >85% were identified.

### Statistical analysis

Data are presented as mean ± standard deviation (SD) from at least three independent experiments. For comparisons among three or more groups, one-way analysis of variance (ANOVA) was performed, followed by Dunnett’s post hoc test for comparisons against the control group. For comparisons between two groups, an unpaired two-tailed Student’s t-test was used. P-values less than 0.05 were considered statistically significant. All analyses were conducted using GraphPad Prism software.

## Supporting information

https://www.ncbi.nlm.nih.gov/geo/query/acc.cgi?acc=GSE312546

## Acknowledgment

This work was supported by the National Natural Science Foundation of China (31571481, 31771585, and 32070799).

## Author Contributions

All authors, including JC (Jingang Cui), FZY (Zhiyu Fan), SD (Suoqi Ding), J.Z. (Jiazhen Zhang), HS (Huihong Shen), SAZ (Syeda Armana Zaidi), and YD (Yongsheng Ding), were involved in the acquisition, analysis, and interpretation of data according to their respective roles. YD conceived and designed the study, drafted the manuscript, and approved the final version for submission.

## Data availability

The data that support the findings of this study are available from the corresponding author upon reasonable request.

