## Supplementary material for "Abnormal shear stress induces ferroptosis in endothelial cells via KLF6 downregulation": https://www.ncbi.nlm.nih.gov/geo/query/acc.cgi?acc=GSE312546

**Raw data of the WB results**

1. **SLC711A in Figure 1F (the shown blots in the red box)**

| **Uncropped blots** | |
| --- | --- |
| **SLC711A**  **(55 kDa）** | 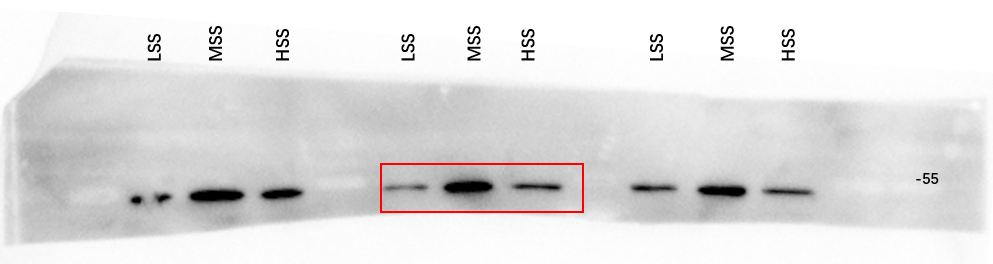 |
| **GAPDH**  **(36 kDa)** | 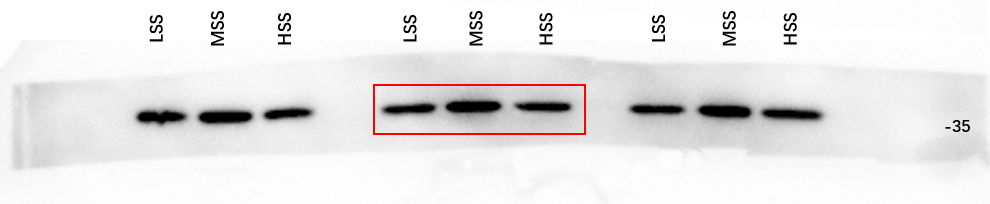 |

2. **BiP in Figure 2G (the shown blots in the red box)**

| **Uncropped blots** | |
| --- | --- |
| **BiP**  **(78 KDa)** | 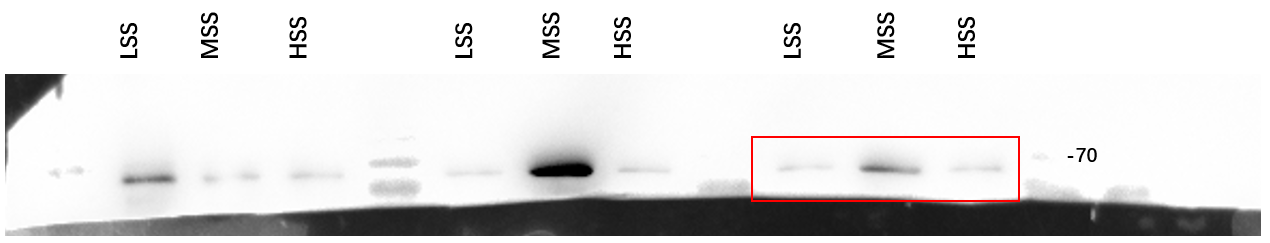  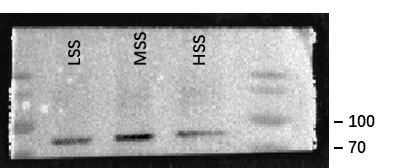 |
| **β-actin**  **(42 kDa)** | 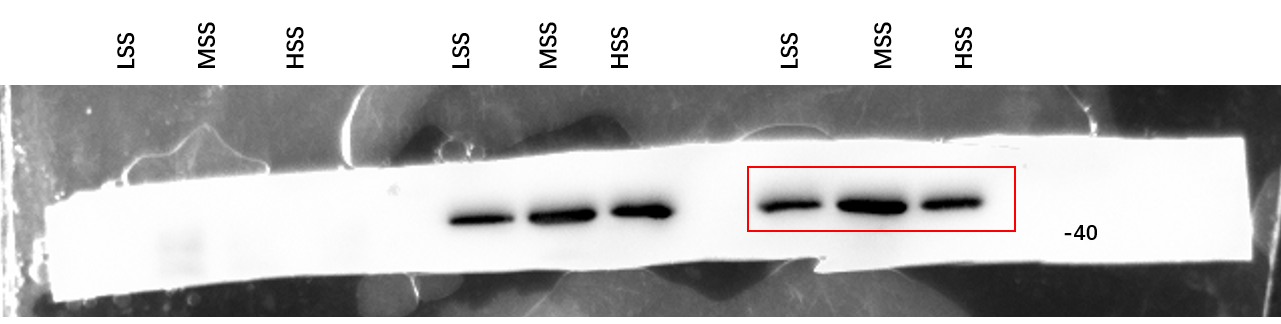  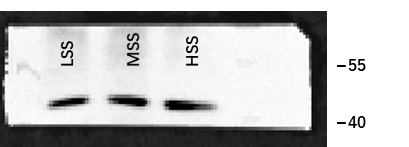 |

3. **PERK in Figure 2G (the shown blots in the red box)**

| **Uncropped blots** | |
| --- | --- |
| **PERK**  **(125 kDa)** | 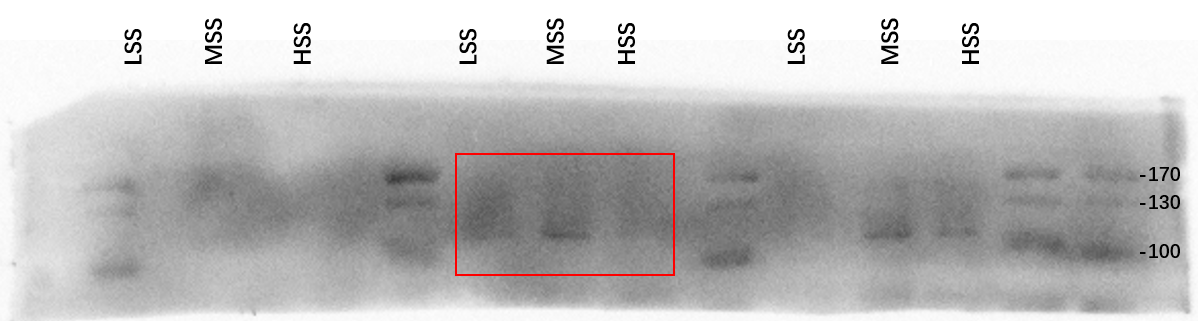  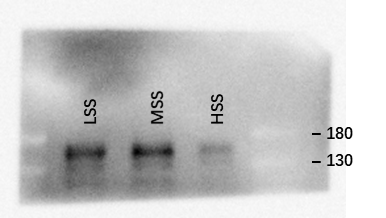 |
| **GAPDH**  **(36 kDa）** | 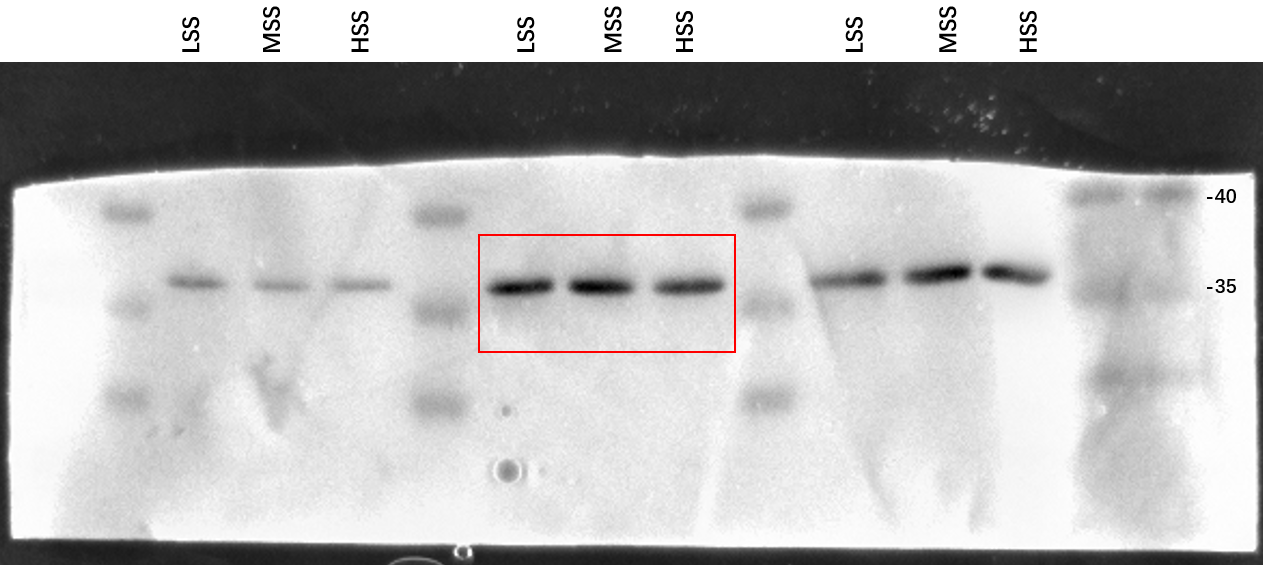  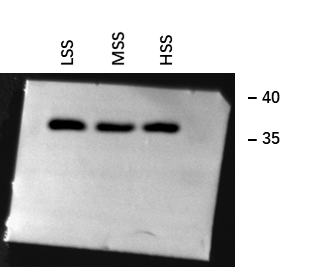 |

4. **MVD in Figure 2G (the shown blots in the red box)**

| **Uncropped blots** | |
| --- | --- |
| **MVD**  **(43 KDa)** | 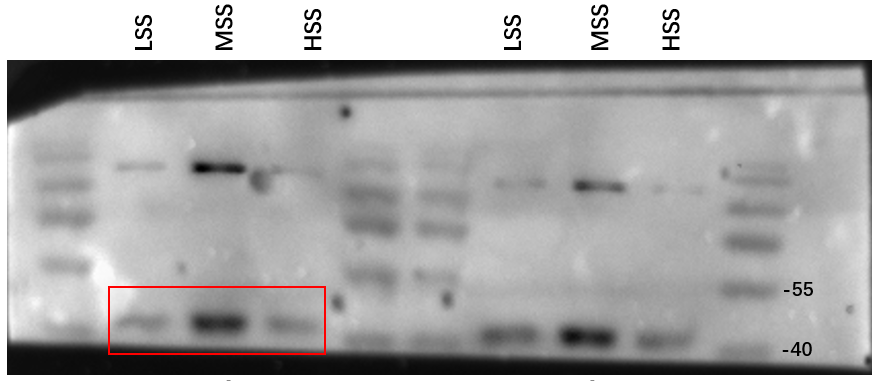  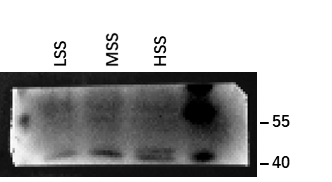 |
| **GAPDH**  **(36 kDa)** | 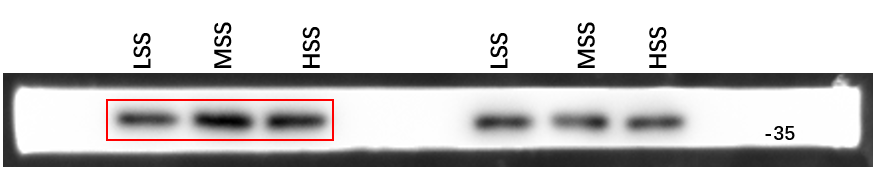  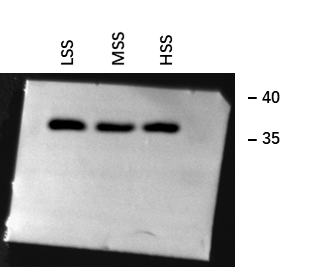 |

5. **IDI1 in Figure 2G (the shown blots in the red box)**

| **Uncropped blots** | |
| --- | --- |
| **IDI1**  **(31 KDa)** | 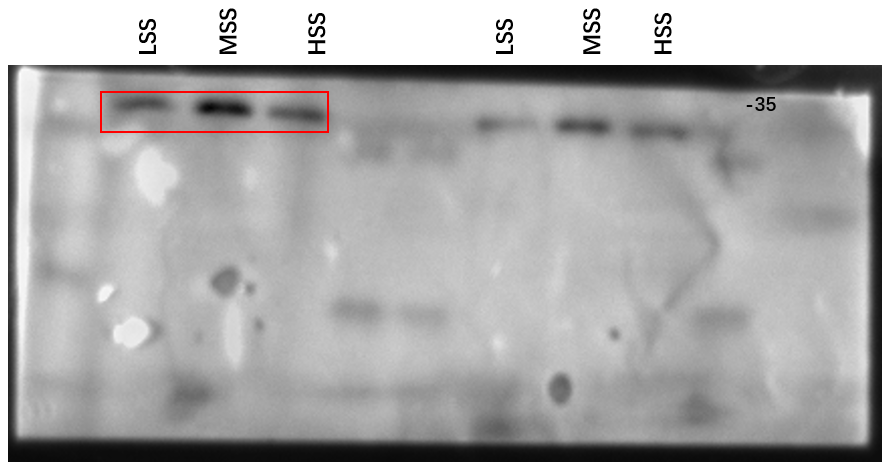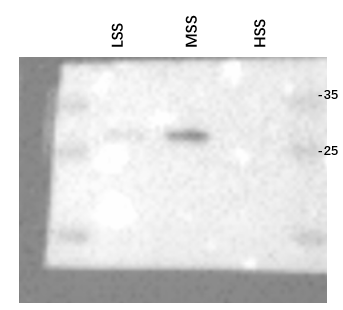 |
| **GAPDH**  **(36 kDa)** | 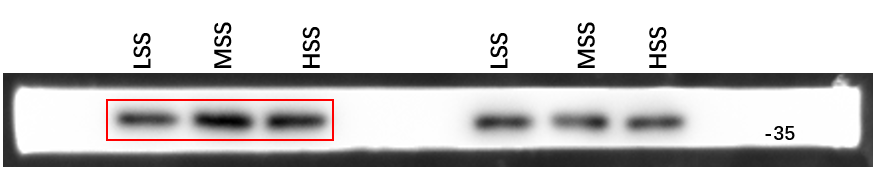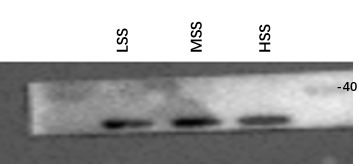 |

6. **KLF6 in Figure 3C (the shown blots in the red box)**

| **Uncropped blots** | |
| --- | --- |
| **KLF6**  **(38 KDa)** | 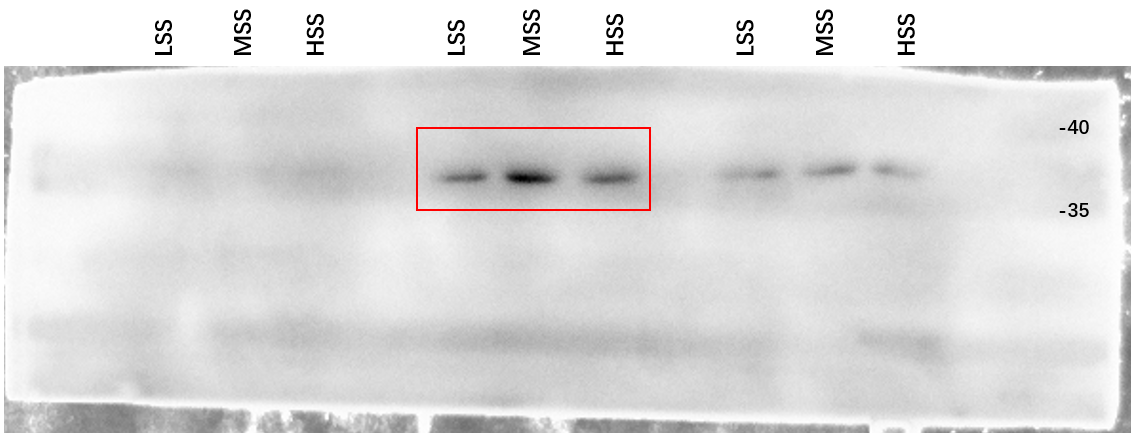  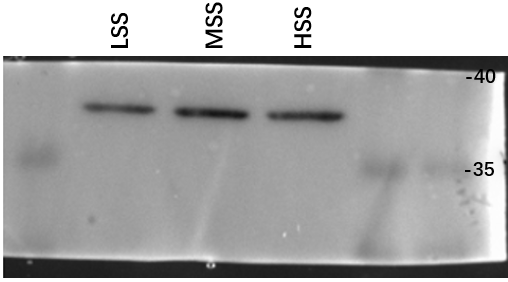 |
| **GAPDH**  **(36 kDa)** | 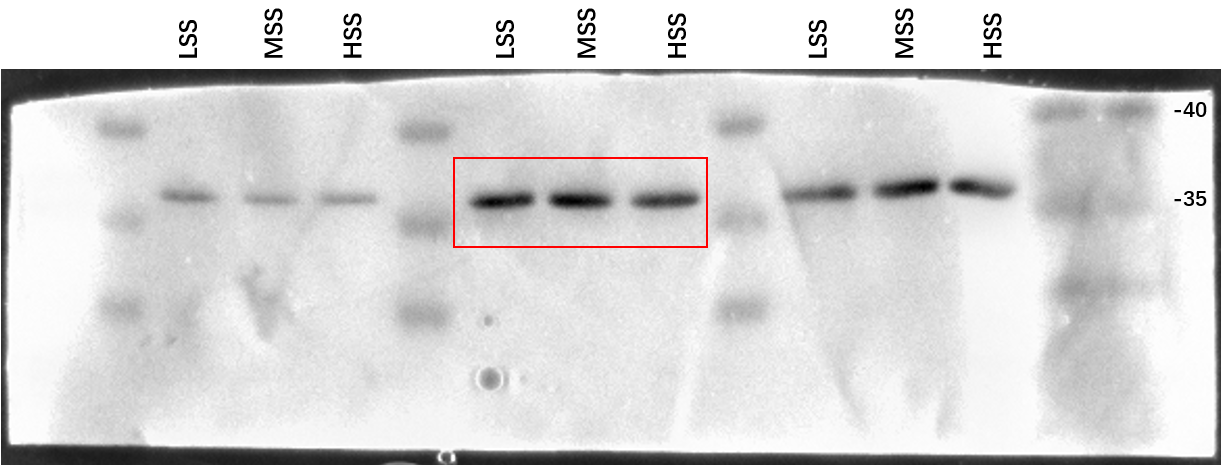  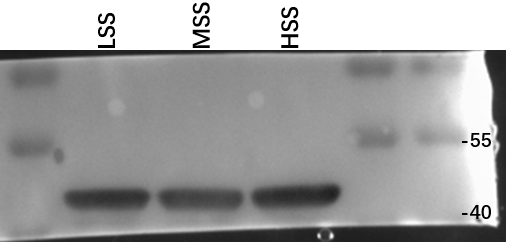  (**β-actin**) |

7. **KLF6 in Figure 3D (the shown blots in the red box)**

| **Uncropped blots** | |
| --- | --- |
| **KLF6**  **(38 KDa)** | 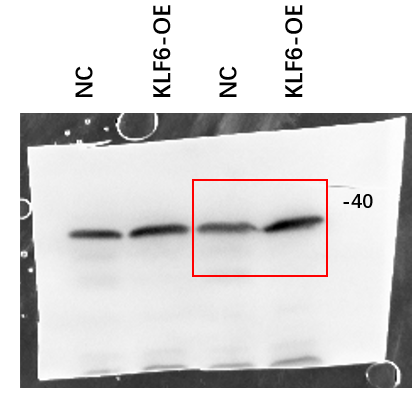  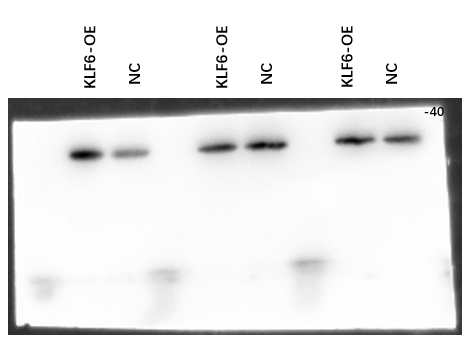 |
| **GAPDH**  **(36 kDa)** | 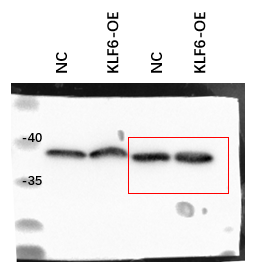  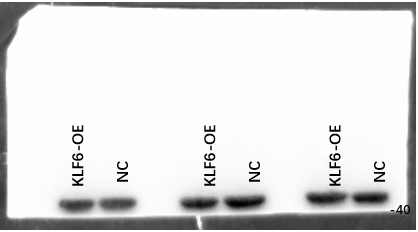 |

8. **KLF6 in Figure 3E (the shown blots in the red box)**

| **Uncropped blots** | |
| --- | --- |
| **KLF6**  **(38 KDa)** | 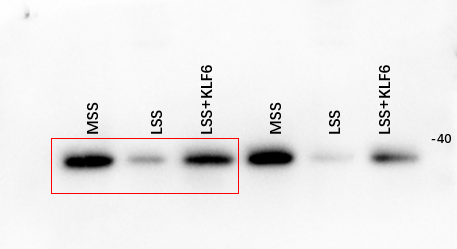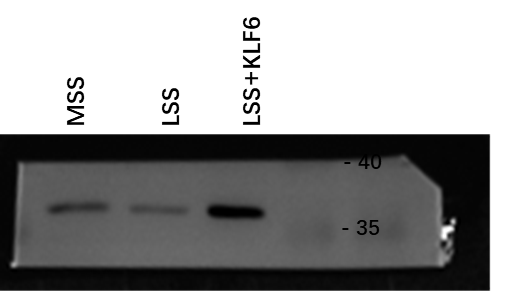 |
| **GAPDH**  **(36 kDa)** | 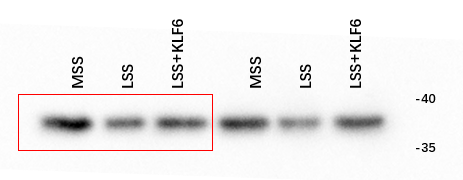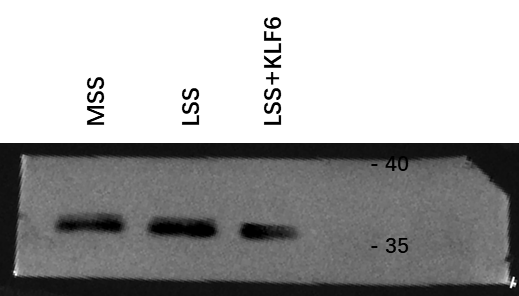 |

9. **PERK in Figure 3E (the shown blots in the red box)**

| **Uncropped blots** | |
| --- | --- |
| **PERK (125 KDa)** | 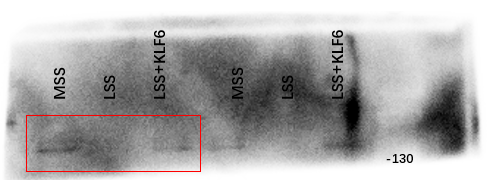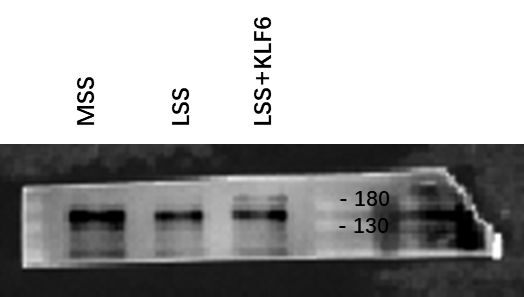 |
| **GAPDH**  **(36 kDa)** | 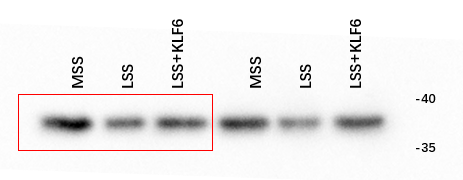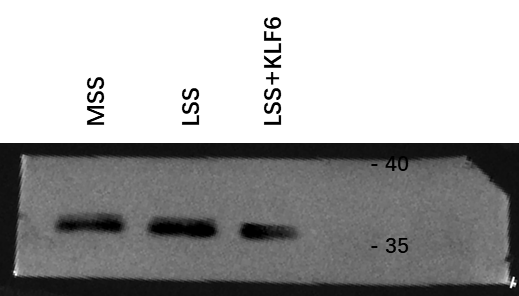 |

10. **BIP in Figure 3E (the shown blots in the red box)**

| **Uncropped blots** | |
| --- | --- |
| **BiP**  **(78 KDa)** | 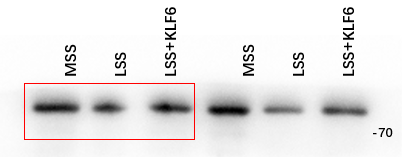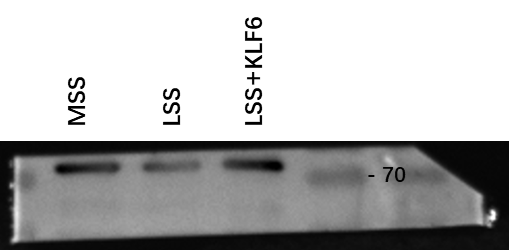 |
| **GAPDH**  **(36 kDa)** | 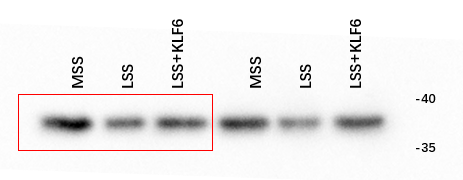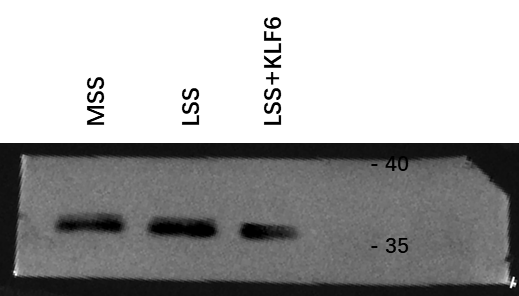 |

11. **MVD in Figure 3E (the shown blots in the red box)**

| **Uncropped blots** | |
| --- | --- |
| **MVD**  **(43 KDa)** | 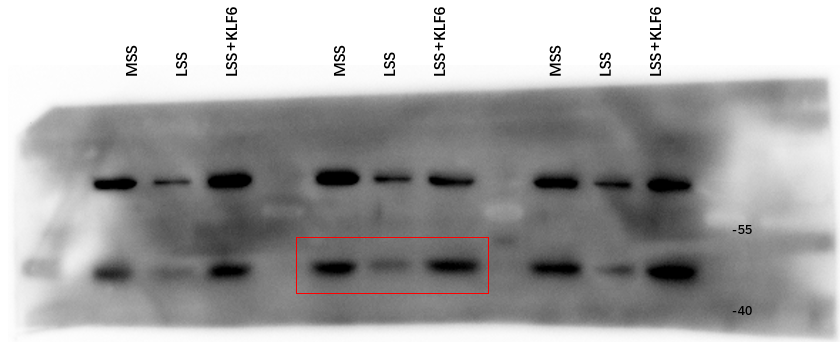 |
| **β-actin**  **(42 kDa)** | 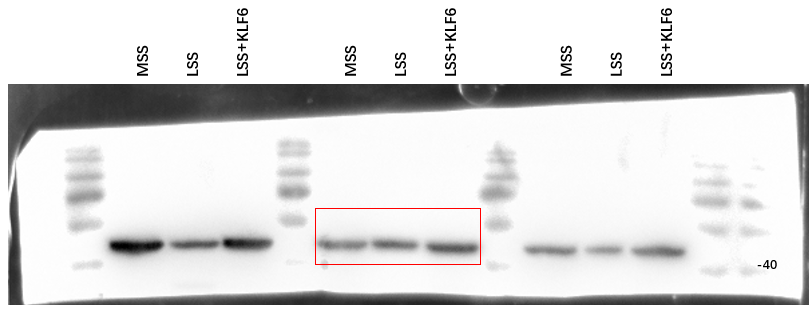 |

12. **IDI1 in Figure 3E (the shown blots in the red box)**

| **Uncropped blots** | |
| --- | --- |
| **IDI1**  **(31 KDa)** | 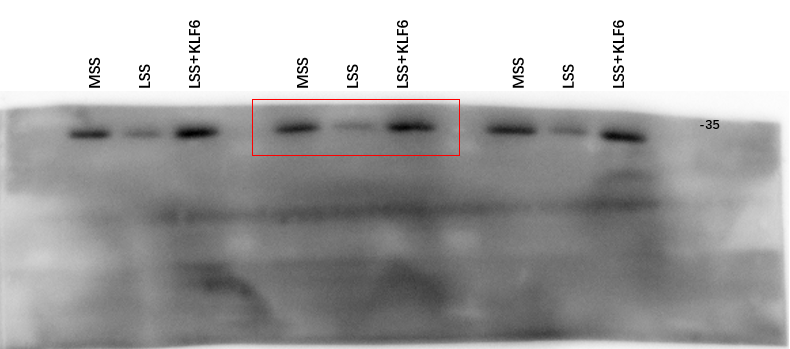 |
| **β-actin**  **(42 kDa)** | 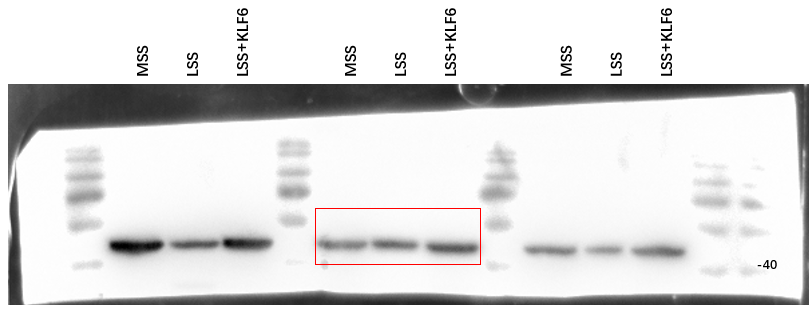 |

13. **KLF6 in Figure 3F (the shown blots in the red box)**

| **Uncropped blots** | |
| --- | --- |
| **KLF6**  **(38 KDa)** | 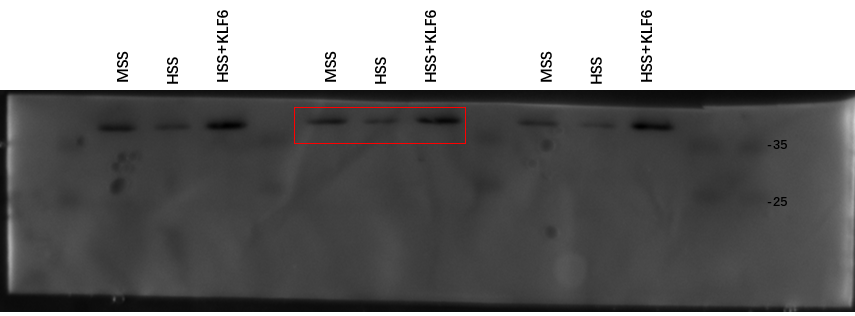 |
| **β-actin**  **(42 kDa)** | 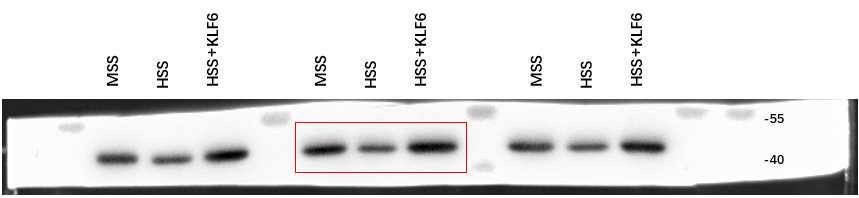 |

14. **PERK in Figure 3F (the shown blots in the red box)**

| **Uncropped blots** | |
| --- | --- |
| **PERK**  **(125 KDa)** | 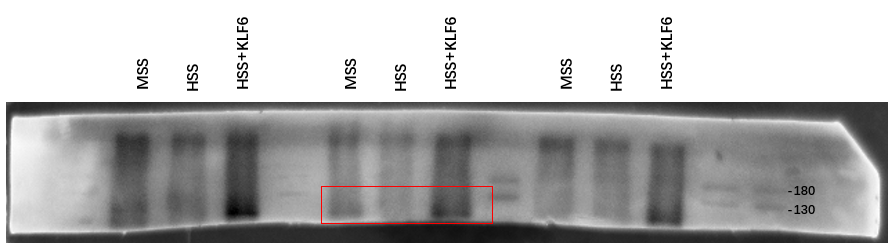 |
| **β-actin**  **(42 kDa)** | 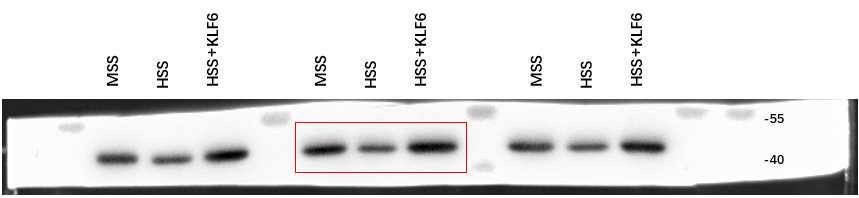 |

15. **BIP in Figure 3F (the shown blots in the red box)**

| **Uncropped blots** | |
| --- | --- |
| **BiP**  **(78 KDa)** | 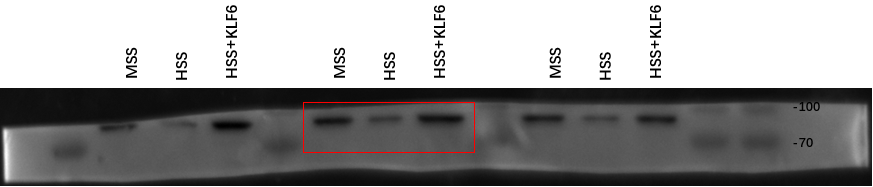 |
| **β-actin**  **(42 kDa)** | 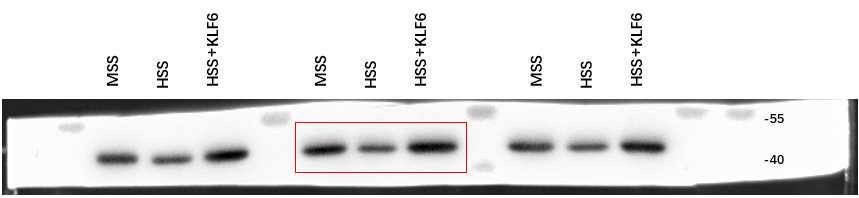 |

16. **MVD in Figure 3F (the shown blots in the red box)**

| **Uncropped blots** | |
| --- | --- |
| **MVD**  **(43 KDa)** | 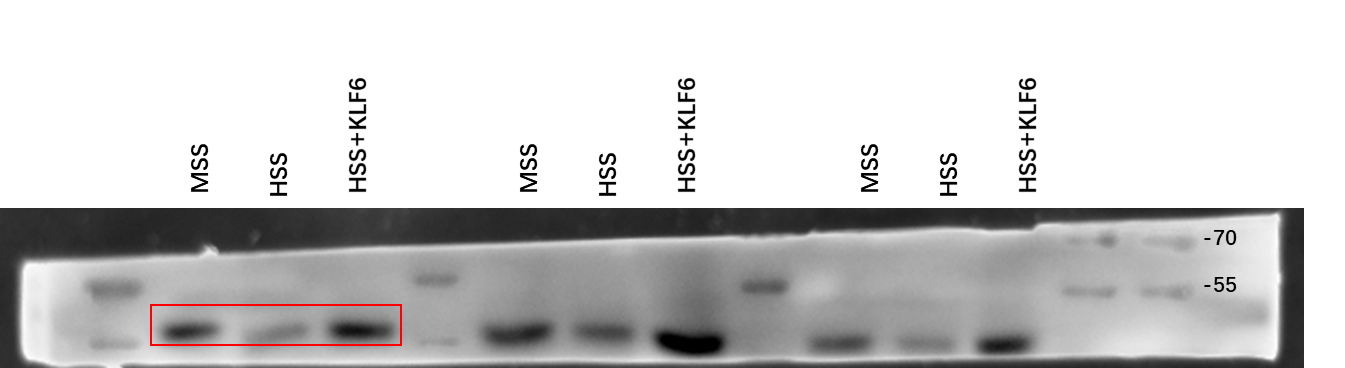 |
| **GAPDH**  **(36 kDa)** | 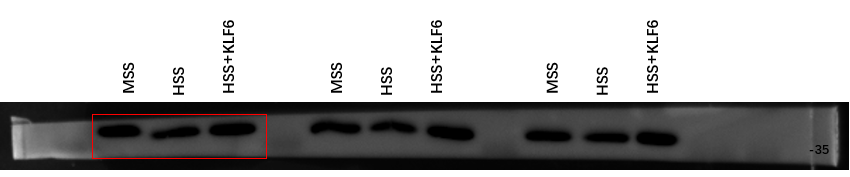 |

17. **IDI1 in Figure 3F (the shown blots in the red box)**

| **Uncropped blots** | |
| --- | --- |
| **IDI1**  **(31 KDa)** | 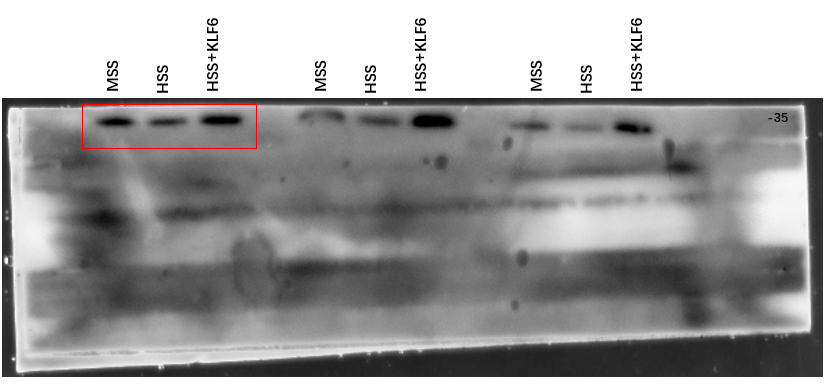 |
| **GAPDH**  **(36 kDa)** | 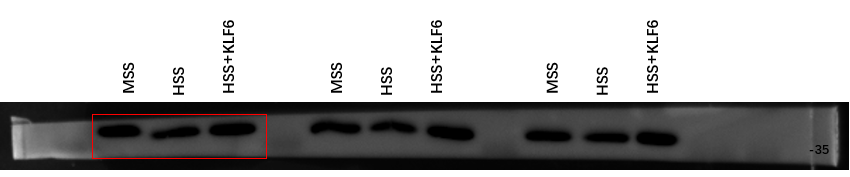 |

18. **SLC7A11 in Figure 4A (the shown blots in the red box)**

| **Uncropped blots** | |
| --- | --- |
| **SLC7A11**  **(55 KDa)** | 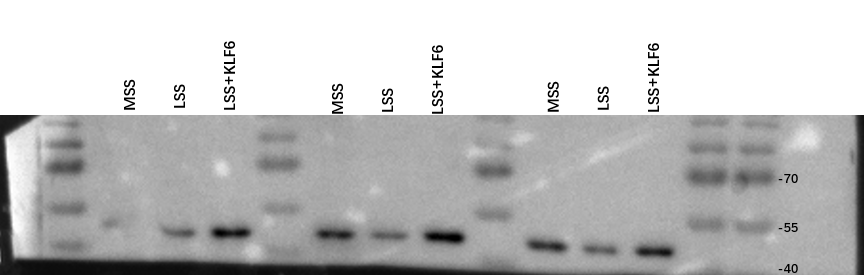  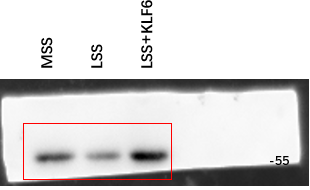 |
| **GAPDH**  **(36 kDa)** | 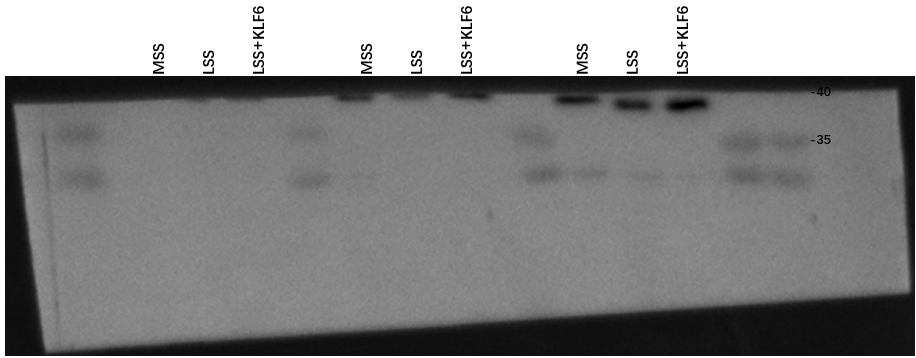  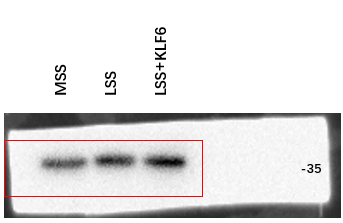 |

19. **SLC7A11 in Figure 4E (the shown blots in the red box)**

| **Uncropped blots** | |
| --- | --- |
| **SLC7A11**  **(55 KDa)** | 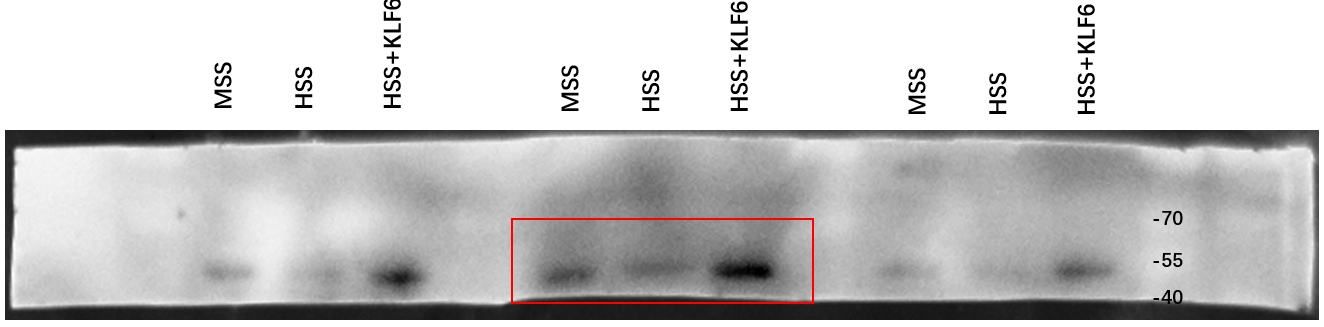 |
| **GAPDH**  **(36 kDa)** | 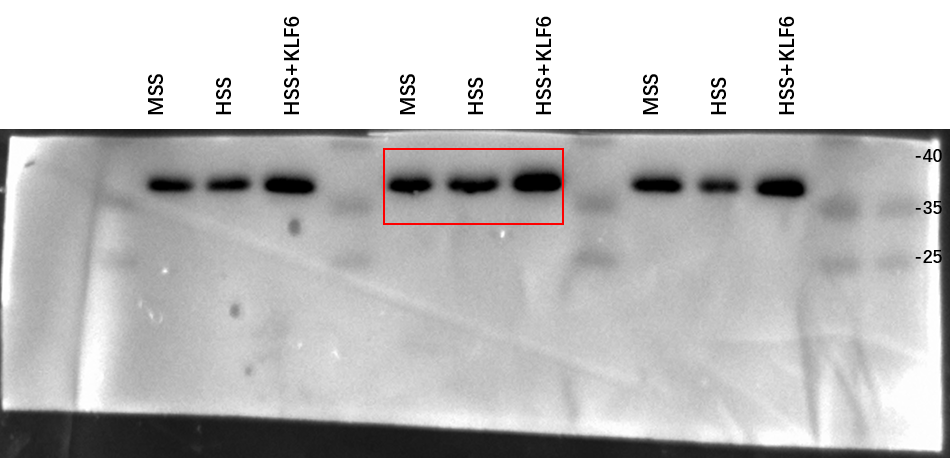 |
